# The inhibitory effects of toothpaste and mouthwash ingredients on the interaction between the SARS-CoV-2 spike protein and ACE2, and the protease activity of TMPRSS2, *in vitro*

**DOI:** 10.1101/2021.03.19.435740

**Authors:** Riho Tateyama-Makino, Mari Abe-Yutori, Taku Iwamoto, Kota Tsutsumi, Motonori Tsuji, Satoru Morishita, Kei Kurita, Yukio Yamamoto, Eiji Nishinaga, Keiichi Tsukinoki

## Abstract

Severe acute respiratory syndrome coronavirus 2 (SARS-CoV-2) enters host cells when the viral spike protein is cleaved by transmembrane protease serine 2 (TMPRSS2) after binding to the host angiotensin-converting enzyme 2 (ACE2). Since ACE2 and TMPRSS2 are expressed in the mucosa of the tongue and gingiva, the oral cavity seems like it is an entry point for SARS-CoV-2. Daily oral care using mouthwash seems to play an important role in preventing SARS-CoV-2 infection. However, the relationship between daily oral care and the mechanisms of virus entry into host cells is unclear. In this study, we evaluated the inhibitory effects of ingredients that are generally contained in toothpaste and mouthwash on the interaction between the spike protein and ACE2 and on the serine protease activity of TMPRSS2 using an enzyme-linked immunosorbent assay and *in vitro* enzyme assay, respectively. Both assays detected inhibitory effects of sodium tetradecene sulfonate, sodium N-lauroyl-N-methyltaurate, sodium N-lauroylsarcosinate, sodium dodecyl sulfate, and copper gluconate. Molecular docking simulations suggested that these ingredients could bind to the inhibitor-binding site of ACE2. In addition, tranexamic acid and 6-aminohexanoic acid, which act as serine protease inhibitors, exerted inhibitory effects on TMPRSS2 protease activity. Further experimental and clinical studies are needed to further elucidate these mechanisms. Our findings support the possibility that toothpaste and mouthwash contain ingredients that inhibit SARS-CoV-2 infection.

## 1 Introduction

Coronavirus disease 2019 (COVID-19), which is caused by severe acute respiratory syndrome coronavirus 2 (SARS-CoV-2), is a serious public health problem worldwide. SARS-CoV-2 mainly infects the upper respiratory tract, which leads to respiratory disease. The viral spike protein mediates SARS-CoV-2 entry into host cells (Lan et al., 2020). To fulfill this function, it is essential that the spike protein bind to angiotensin-converting enzyme 2 (ACE2) on the host cells (Lan et al., 2020). The transmembrane protease serine 2 (TMPRSS2) of the host is an essential factor of SARS-CoV-2 entry into host cells (Hoffmann et al., 2020b). TMPRSS2 cleavages the spike protein bound to ACE2, and the cleaved spike protein leads to the fusion of viral and host cell membranes (Hoffmann et al., 2020b).

The oral cavity is an important entry point for pathogens. Xu et al. demonstrated that ACE2 was expressed in oral mucosa, especially the dorsal tongue (Xu et al., 2020). Sakaguchi et al. and Huang et al. showed that ACE2 and TMPRSS2 are expressed in the mucosa of the tongue, gingiva, and salivary gland epithelial cells (Huang et al., 2020; Sakaguchi et al., 2020). These findings suggest that multiple oral epithelial cells serve as an entry point for SARS-CoV-2. Although whether the activities of ACE2 and TMPRSS2 on the tongue are associated with virus infection is unclear, taste impairment has been recognized as a symptom of COVID-19 (Yan et al., 2020). Huang et al. suggested that SARS-CoV-2 infects the salivary glands and oral mucosa and implicates saliva in viral transmission (Huang et al., 2020). These findings support that the oral cavity is an entry point for SARS-CoV-2. In a critical review, Carrouel et al. suggested that some ingredients in antiseptic mouthwash demonstrate antiviral properties (Carrouel et al., 2020). Seneviratne et al. demonstrated that mouthwashes containing povidone iodine or cetylpyridinium chloride exhibit the potential to decrease SARS-CoV-2 viral load in saliva (Seneviratne et al., 2020). Therefore, daily oral care may play an important role in preventing SARS-CoV-2 infection, but the relationship between daily oral care and the mechanisms of virus entry into host cells are unclear.

This study aimed to investigate the effects of general ingredients contained in commercially available toothpastes and mouthwashes on the mechanisms of SARS-CoV-2 entry into host cells. Specifically, we evaluated the inhibitory effects of these ingredients on the interaction between the spike protein of SARS-CoV-2 and human ACE2, as well as the protease activity of TMPRSS2, *in vitro*. We also performed docking simulations for these target proteins to support the experimental results.

## 2 Materials and Methods

### 2.1 Materials

The test ingredients (Table 1) were purchased from FUJIFILM Wako Pure Chemical Corporation, Nikko Chemicals Co., Ltd., and Lion Specialty Chemicals Co., Ltd. Each stock solution of test ingredients was diluted with phosphate-buffered saline (PBS) at a concentration of 5% (w/w). Recombinant human TMPRSS2 (N-terminus 6xHis, aa106-492) was purchased from LifeSpan Biosciences (Seattle, WA, USA). Boc-Gln-Ala-Arg-MCA were purchased from Peptide Institute Inc. (Osaka, Japan).

**Table 1.**
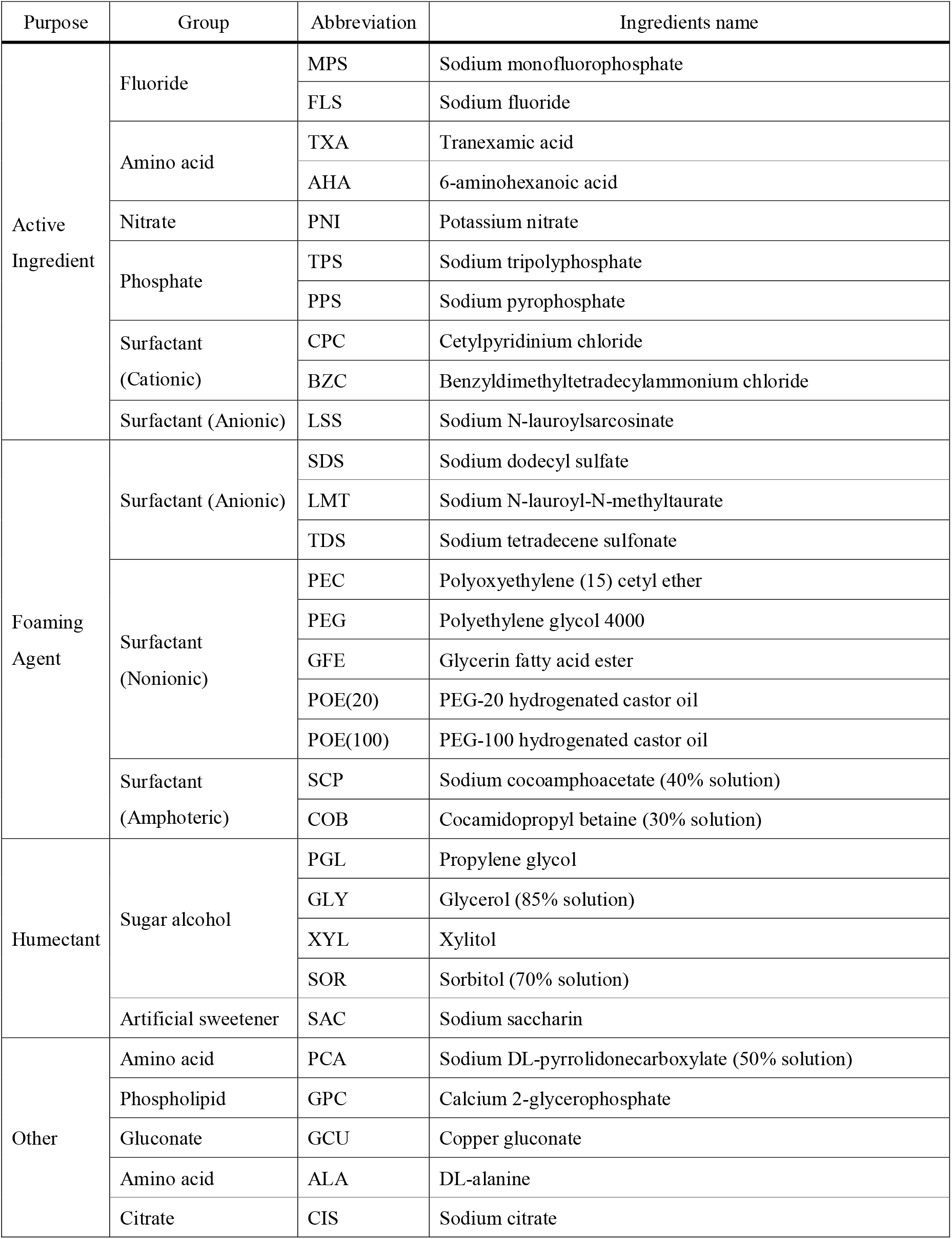
List of test ingredients used for *in vitro* screening

### 2.2 *In vitro* assay of the interaction between receptor-binding domain of spike protein and ACE2

The interaction between the receptor-binding domain (RBD) of the SARS-CoV-2 spike protein and ACE2 was estimated using Spike S1 (SARS-CoV-2): ACE2 Inhibitor Screening Colorimetric Assay Kit (BPS Bioscience, San Diego, CA, USA). The absorbance value (AV) at 450 nm was read using Infinite 200 Pro (Tecan Japan, Kanagawa, Japan). All assays were performed, following the manufacturer’s instructions. The inhibitory rate was calculated as follows: Inhibition (%) = (AV of control – AV of ingredients treatment) / AV of control × 100. The half-maximal inhibitory concentration (IC50) was calculated using the DRC package in R software program (v3.6.1).

### 2.3 *In vitro* assay of TMPRSS2 serine protease activity

The serine protease activity of TMPRSS2 was estimated as previously described by Hoffmann et al and Shrimp et al. (Hoffmann et al., 2020a; Shrimp et al., 2020). Recombinant human TMPRSS2 (4 μg/mL final concentration) diluted with an assay buffer (50 mM Tris-HCl pH 8.0, 154 mM NaCl) and test ingredients were added to the 384 well black plate (Greiner Bio-One Japan, Tokyo, Japan). Then, Boc-Gln-Ala-Arg-MCA (10 μM final concentration) diluted with an assay buffer containing dimethyl sulfoxide (DMSO; 0.1% (w/w) final concentration) was added to induce an enzyme reaction. After incubation at room temperature for 1 hr, the fluorescence intensity (FI) of fluorescent hydrolysate of Boc-Gln-Ala-Arg-MCA (7-amino-4-methylcoumarin) were read using SpectraMax M5 plate reader (Molecular Devices, San Jose, CA, USA) with excitation of 380 nm and emission of 460 nm. The inhibitory rate of ingredients was calculated as follows: Inhibition (%) = (FI of control – FI of treatment) / FI of control × 100. The IC50 was calculated using the DRC package in the R software program (v3.6.1).

### 2.4 Preparation of 3D structures of the target proteins for docking simulations

We prepared a human ACE2 structure that is suitable for docking simulations using a crystal structure (PDB ID: 1R4L (Towler et al., 2004); resolution 3.00 Å). Structural refinement was performed using Homology Modeling Professional for HyperChem (HMHC) software (Tsuji, 2007; Tsuji et al., 2015). Conversely, an X-ray crystal structure of human TMPRSS2 has never been resolved. According to the reference (Rahman et al., 2020), a human TMPRSS2 model that is suitable for docking simulations was prepared by the SWISS-MODEL (Waterhouse et al., 2018), using a template (PDB ID: 5CE1; human serine protease complexed with inhibitor). The inhibitor of 5CE1 was extracted for the prepared model using HMHC. Then, the N- and C-terminals of both structures were treated as zwitterions; aspartic and glutamic acid residues were treated as anions; and lysine, arginine, and histidine residues were treated as cations under the physiological conditions. Other small molecules except for the inhibitor, Zn^2+^ ion, Cl^-^ ion, and crystal waters were removed. The detailed preparation procedure was described previously in detail (Tsuji, 2020).

A 3D structure of the seven test compounds was downloaded from the PubChem website in an SDF file format. These structures were converted into individual PDBQT files under the physiological condition of pH = 7.4 using a Docking Study with HyperChem (DSHC) software (Tsuji, 2007; Tsuji et al., 2015).

### 2.5 Validation of the docking studies with crystal inhibitors and subsequent docking simulations for the test ingredients

The above prepared target protein structures and inhibitor structures were converted to the PDBQT files using DSHC software. Configuration files for performing AutoDock Vina (Trott and Olson, 2010) docking simulations were prepared based on the coordinates of the corresponding crystal inhibitor using the AutoDock Vina In Silico Screening Interface of DSHC. The exhaustiveness value was set to 100, and the top nine docking modes were maximally output. This docking condition was also used for subsequent docking simulations of the individual test compounds (Tsuji, 2020).

## 3 Results

### 3.1 Inhibitory effects of toothpaste and mouthwash ingredients on the interaction between the spike protein of SARS-CoV-2 and ACE2

To understand the effect of commercially available toothpaste and mouthwash on the SARS-CoV-2 entry mechanisms into host cells, we evaluated 30 general ingredients (Table 1). We first screened the 30 ingredients for inhibitory effects on the interaction between the spike protein of SARS-CoV-2 and ACE2 using enzyme-linked immunosorbent assay. When analyzed at 1% (w/w), 18 ingredients showed a > 0% inhibitory rate (Fig. 1). Five ingredients (TDS, LMT, LSS, GCU, and SDS) exhibited greater than 50% inhibitory rates. The inhibitory rates of TDS, LMT, LSS, GCU, and SDS at 1% (w/w) were 96.7%, 95.9%, 94.9%, 94.5%, and 92.9%, respectively. To clarify the details of the inhibitory effects and to determine the IC50 values, we assessed the dose-effect relationship of these five ingredients. TDS, LMT, and LSS ranged from 0.00005% to 1% (w/w); SDS ranged from 0.0005% to 1% (w/w); and GCU ranged from 0.005% to 1% (w/w) (Fig. 2). TDS, LMT, LSS, GCU, and SDS showed inhibitory activities against the interaction between the RBD of the spike protein and ACE2 with IC50 values of 0.009, 0.022, 0.043, 0.097, and 0.005% (w/w), respectively (Fig. 2).

**Figure 1.**
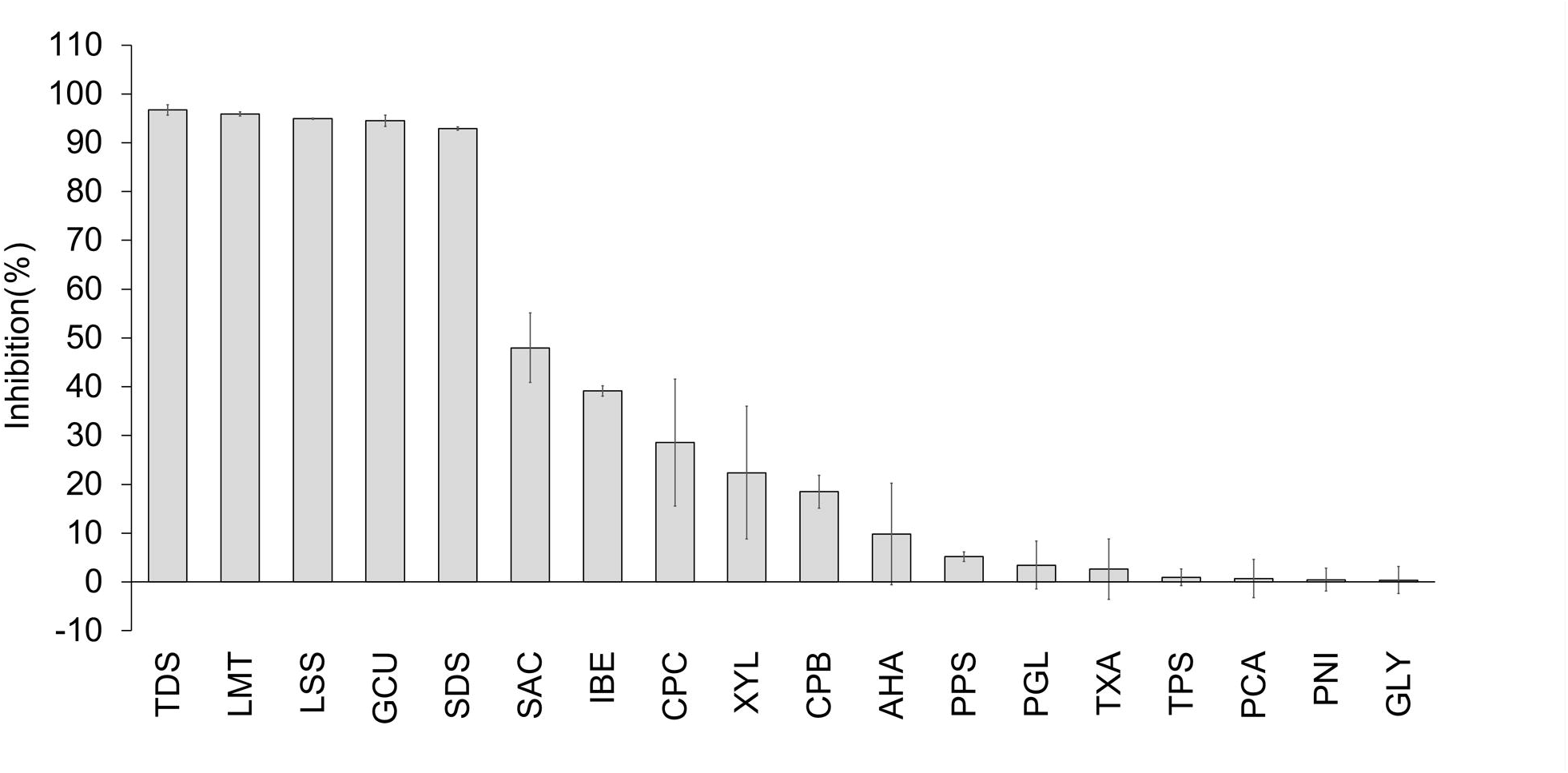
Screening of toothpaste and mouthwash ingredients exhibited inhibitory effects on the interaction between the spike protein RBD of SARS-CoV-2 and ACE2. Eighteen toothpaste and mouthwash ingredients at 1% (w/w) exhibited inhibitory effects interaction between the spike protein of SARS-CoV-2 and ACE2 *in vitro*. The interaction between the spike protein RBD of SARS-CoV-2 and ACE2 was evaluated by measuring OD at 450 nm using a microplate reader. Data are expressed as the mean ± SD (n = 3).

**Figure 2.**
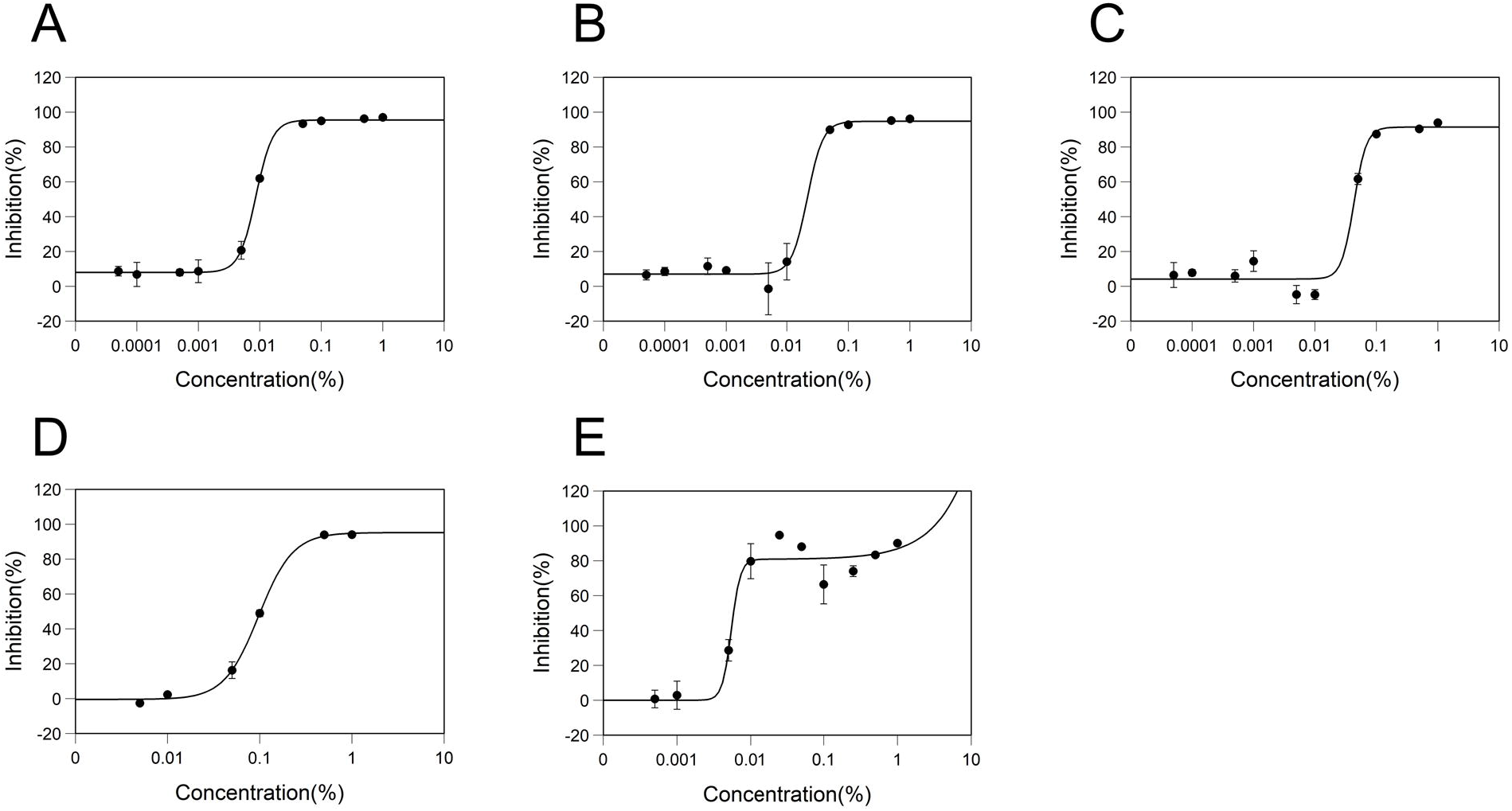
Effect of ingredient concentration on inhibitory effects of the interaction between the spike protein of SARS-CoV-2 and ACE2. The dose-response inhibitory effects of interaction between spike protein of SARS-CoV-2 and ACE2 for (A) Sodium tetradecene sulfonate, (B) Sodium N-lauroyl-N-methyltaurate, (C) Sodium N-lauroylsarcosinate, (D) Copper gluconate, and (E) Sodium dodecyl sulfate are shown. The data were plotted and modeled by a four-parameter Log-logistic fit (A-D) and a four-parameter Brain-Cousens fit (E) to determine the 50% inhibitory concentration (IC50) value. All data points were expressed as the mean ± SD (n = 3)

### 3.2 Inhibitory effects of toothpaste and mouthwash ingredients on TMPRSS2 protease activity

We next screened the 30 ingredients (Table 1) for inhibitory effects on the serine protease activity of TMPRSS2. The TMPRSS2 activity test was carried out using recombinant human TMPRSS2 and a synthetic peptide substrate (Hoffmann et al., 2020a; Shrimp et al., 2020). When analyzed at 1 % (w/w), 20 ingredients showed a > 0% inhibitory rate. CPC and SAC clearly exhibited false-positive reactions in an assay in which the substrate, Boc-Gln-Ala-Arg-MCA, was replaced with 7-amino-4-methylcoumarin; therefore, CPC and SAC were excluded from this evaluation (data not shown). Therefore, in this study, 18 ingredients with an inhibition rate > 0% (Fig. 3) were selected for this study. Seven ingredients (SDS, TDS, GCU, TXA, LMT, LSS, and AHA) exhibited inhibitory rates above 50%. The inhibitory rate of SDS, TDS, GCU, TXA, LMT, LSS, and AHA at 1% (w/w) was 99.9%, 97.5%, 97.2%, 92.0%, 88.6%, 87.6%, and 71.7%, respectively. To clarify the details of the inhibitory effects and determine the IC50 values, we assessed the dose-effect relationship of these seven ingredients using IC50 values from 0.0005% to 1% (w/w) (Fig. 4). SDS, TDS, GCU, TXA, LMT, LSS, and AHA showed inhibitory activities against the protease activity of TMPRSS2 with IC50 values of 0.014, 0.018, 0.411, 0.054, 0.098, 0.102, and 0.449 % (w/w), respectively (Fig. 4).

**Figure 3.**
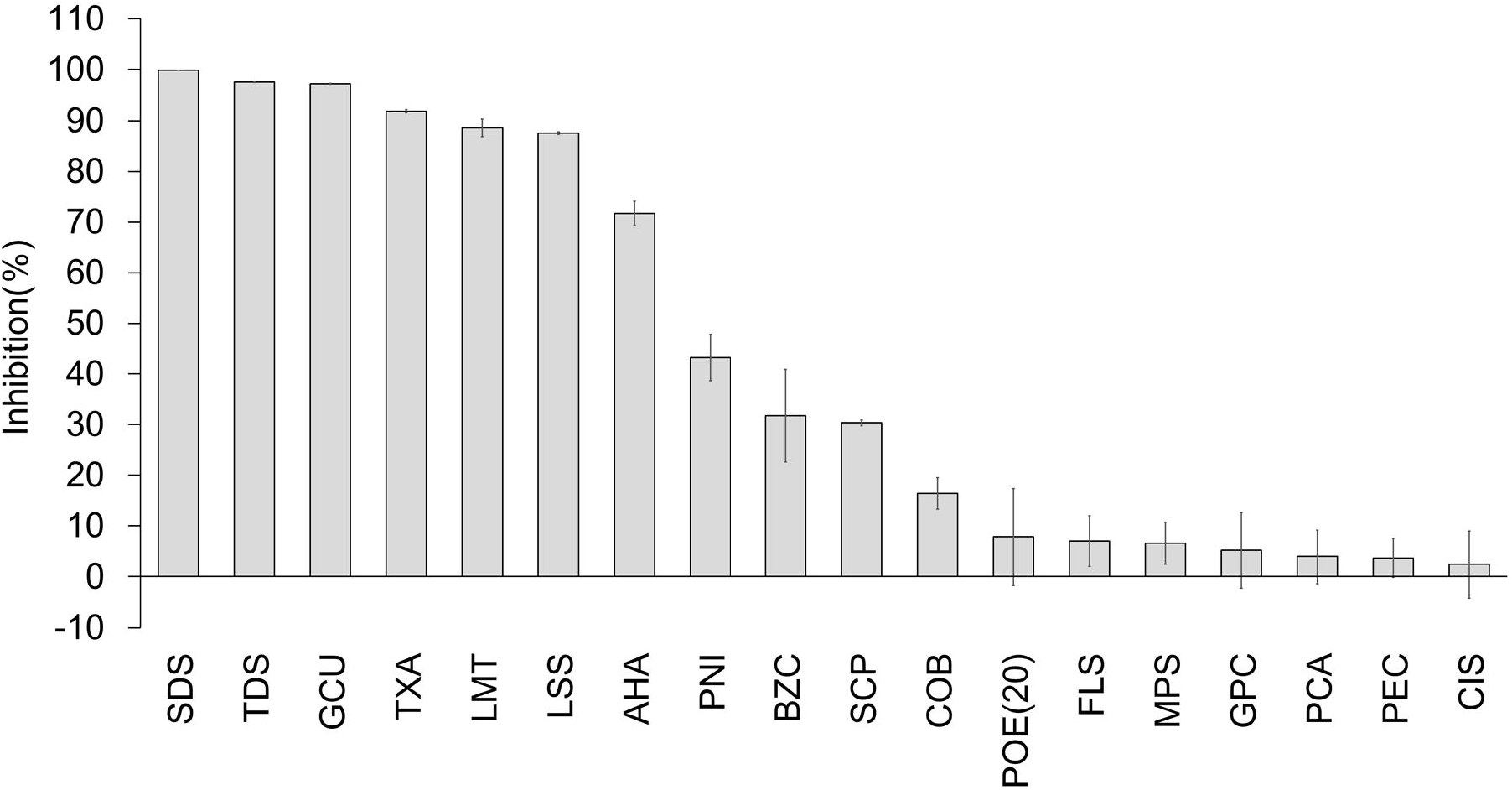
Screening of toothpaste and mouthwash ingredients exhibited inhibitory effects on the serine protease activity of TMPRSS2. Eighteen toothpaste and mouthwash ingredients at 1% (w/w) exhibited inhibitory effects on the TMPRSS2 serine protease activity. TMPRSS2 cleaved Boc-Gln-Ala-Arg-MCA as the substrate and produced the potent fluorophore, AMC (7-amino-4-methylcoumarin). Values were normalized against the intensity of the absence of test ingredients. Data are expressed as the mean ± SD (n = 3).

**Figure 4.**
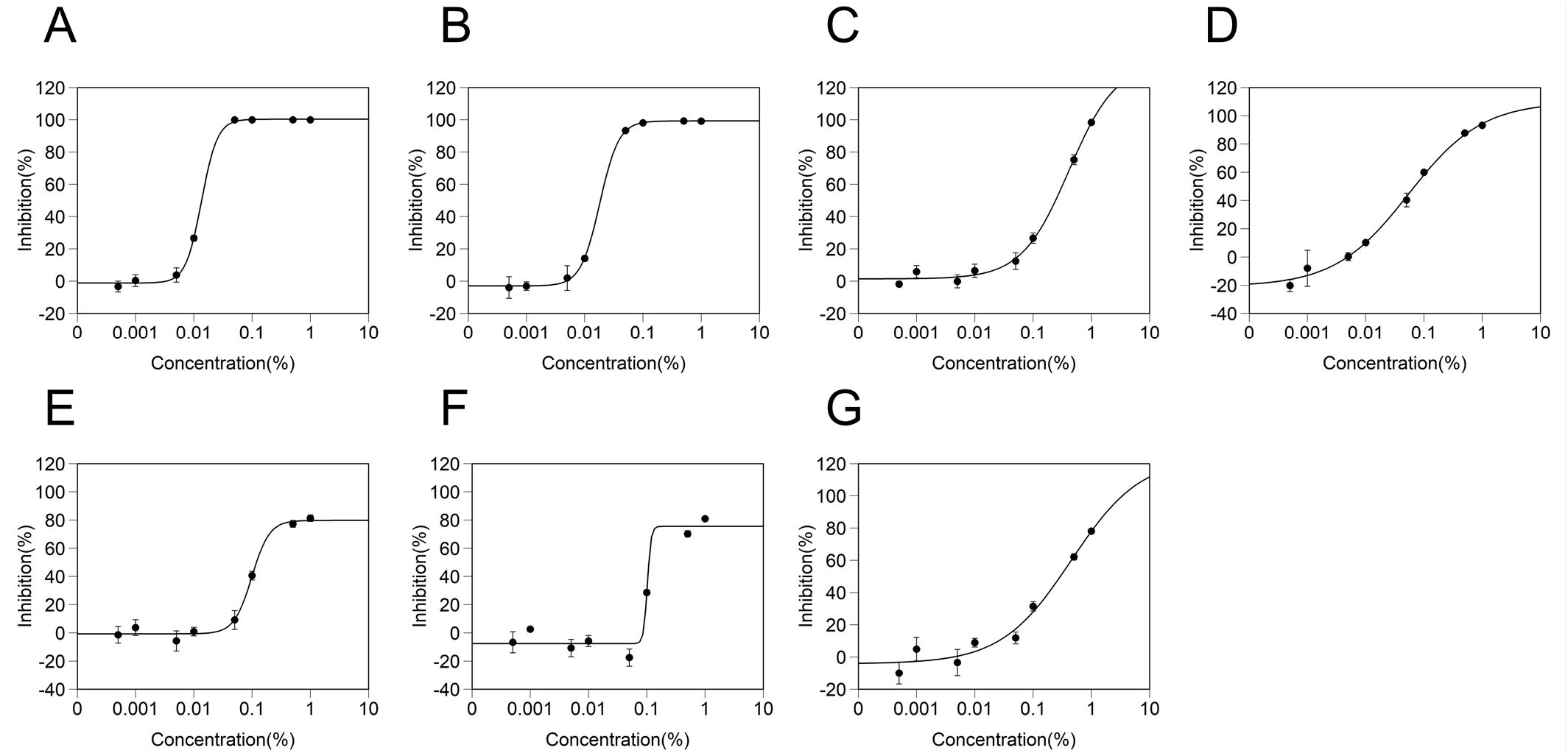
Effect of ingredient concentration on inhibitory effects of serine protease activity of TMPRSS2. The dose-response inhibitory effects of TMPRSS2 serine protease activity by A) Sodium dodecyl sulfate, B) Sodium tetradecene sulfonate, C) Copper gluconate, D) Tranexamic acid, E) Sodium N-lauroyl-N-methyltaurate, F) Sodium N-lauroylsarcosinate, and G) 6-aminohexanoic acid are shown. The data were plotted and modeled by a four-parameter Log-logistic fit to determine the 50% inhibitory concentration (IC50) value. All data points expressed as the mean ± SD (n = 3).

### 3.3 Docking simulations of toothpaste, mouthwash, and their ingredients in the human ACE2 and human TMPRSS2 model

To support the *in vitro* assay, we performed a molecular docking simulation. In the molecular docking study, we used an X-ray crystal structure (PDB ID: 1R4L) for human ACE2 in closed conformation with a synthetic inhibitor

((*S,S*)-2-{1-carboxy-2-[3-(3,5-dichloro-benzyl)-3*H*-imidazol-4-yl]-ethylamino}-4-methyl-pentanoic acid) (Towler et al., 2004). The closed conformation of ACE2 prevented binding to the SARS-CoV-2 spike protein (Reznikov et al., 2021). To determine the docking conditions, we performed a re-docking study using the synthetic inhibitor complexed in the crystal structure. As for the results, the most stable docking mode obtained from the re-docking study reproduced the crystal structure shown in Fig. 5 A, B. The X-ray crystal structure of human TMPRSS2 has not been resolved to date. We performed docking simulations using the homology model obtained from SWISS-MODEL using a template (PDB ID: 5CE1; human serine protease complexed with an inhibitor, 2-[6-(1-hydroxycyclohexyl)pyridin-2-yl]-1*H*-indole-5-carboximidamide) as previously described (Rahman et al., 2020). Although the previous study (Rahman et al., 2020) searched the binding site of the TMPRSS2 inhibitor camostat mesylate in the model using Molecular Operating Environment (MOE) software, we performed docking simulations at the crystal inhibitor-binding site. The re-docking study reproduced a crystal structure of the inhibitor, as shown in Fig. 5 C, D.

**Figure 5.**
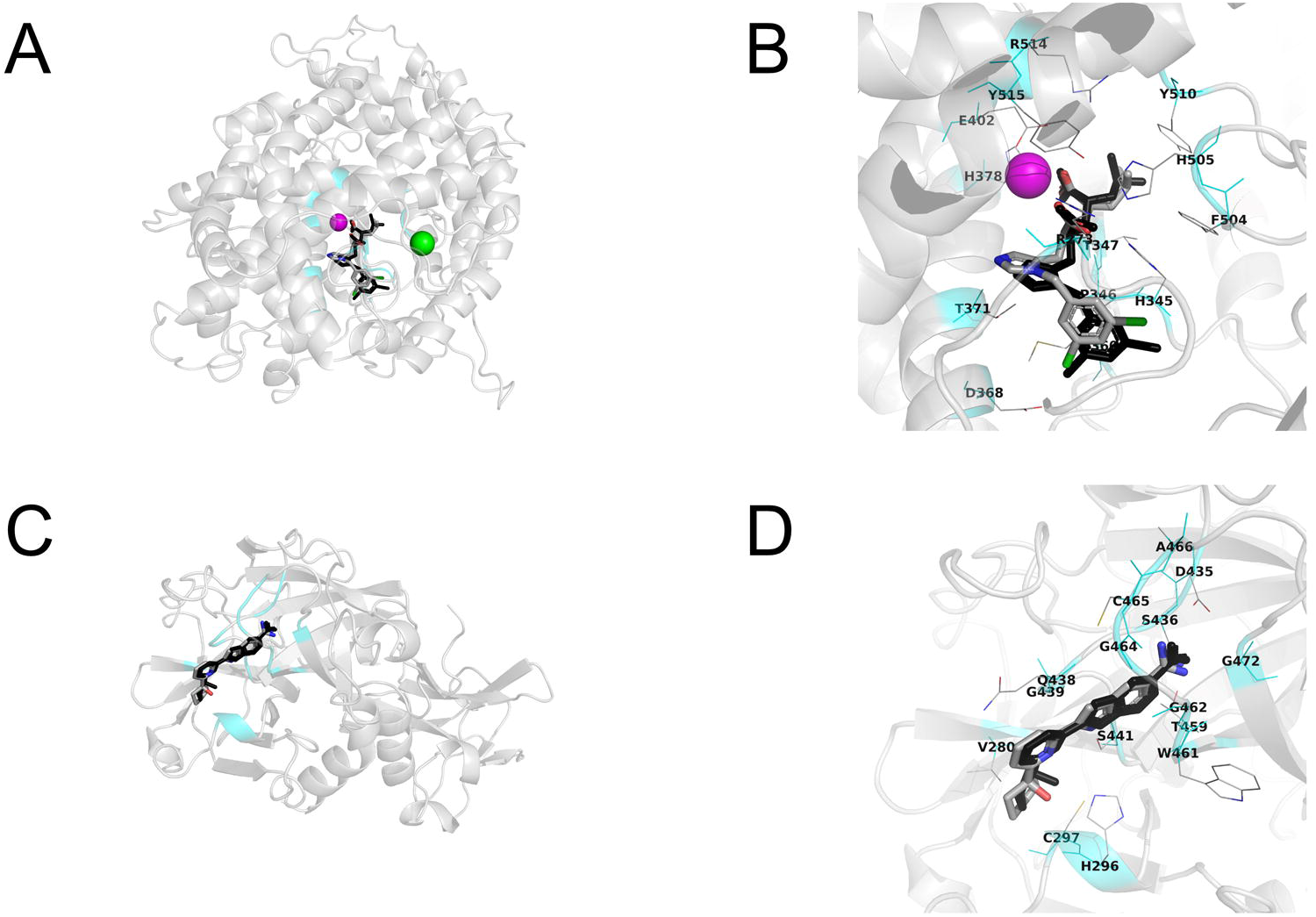
Re-docking studies for human ACE2 and human TMPRSS2 model with their crystal inhibitors. A, B) Human ACE2. C, D) Human TMPRSS2 model. The most stable docking mode of crystal inhibitor shows in CPK color using tubes. The original crystal structure of the inhibitor is represented by black tubes. A magenta sphere shows Zn^2+^ ion and a green sphere shows the Cl^-^ ion. The inhibitor-binding site located at 3 Å from all heavy atoms of the crystal inhibitor is shown in cyan color. Amino acid residues located at 3 Å from the inhibitor are shown using thin tubes with label. Hydrogen atoms are neglected.

Based on the conditions confirmed above, the docking simulations of the seven test ingredients (Fig. 2, 4, Table 2), whose IC_50_ values were estimated using *in vitro* assay, were performed using AutoDock Vina program by targeting the proteins. Incidentally, ingredients used for docking simulations removed counter ions from the test ingredients (Table 2). Figure 6 shows the most stable docking mode, except for 6-aminohexanoic acid, tranexamic acid, and gluconic acid used for docking with the human TMPRSS2 model, which were obtained from AutoDock Vina docking simulations. For the human TMPRSS2 model, reasonable docking modes of 6-aminohexanoic acid, tranexamic acid, and gluconic acid were observed within the top nine docking mode scores. Tables 3 and 4 show the AutoDock Vina scores (empirical binding free energy: *ΔG*_bind_ (kcal/mol)) of these ingredients. The obtained AutoDock Vina scores and docking modes indicated that *N*-lauroyl-*N*-methyltaurine, *N*-lauroylsarcosine, gluconic acid, dodecyl sulfate, and (*E*)-tetradec-1-ene-1-sulfonic acid could weakly bind to human ACE2 at the inhibitor-binding site (Table 3, Fig. 6 A) (Kwofie et al., 2019; Zhong et al., 2019). Additionally, gluconic acid and tranexamic acid could weakly bind to human TMPRSS2 at the inhibitor-binding site (Table 4, Fig. 6 B) (Zhong et al., 2019; Andleeb et al., 2020). These results supported the results of the *in vitro* assay.

**Table 2.** Test ingredients subjected to docking simulations

| Ingredients name | Abbreviation | Ingredients used for docking simulations |  |
| --- | --- | --- | --- |
|  |  | Name | Pubchem CID |
| 6-aminohexanoic acid | AHA | 6-aminocaproic acid | 564 |
| Tranexamic acid | TXA | Tranexamic acid | 5526 |
| Sodium N-lauroylsarcosinate | LSS | N-Lauroylsarcosine | 7348 |
| Sodium dodecyl sulfate | SDS | Dodecyl sulfate | 8778 |
| Copper gluconate | GCU | Gluconic acid | 10690 |
| Sodium N-lauroyl-N-methyltaurine | LMT | N-Lauroyl-N-methyltaurine | 61353 |
| Sodium tetradecene sulfonate | TDS | (E)-Tetradec-1-ene-1-sulfonic acid | 6437821 |

**Table 3.** Vina score of test ingredients for human ACE2

| Ingredients |  | Vina Score<br>(kcal/mol) |
| --- | --- | --- |
| Name | Pubchem CID |  |
| Inhibitor | - | -9.1 |
| N-Lauroyl-N-methyltaurine | 61353 | -6.6 |
| N-Lauroylsarcosine | 7348 | -6.3 |
| Gluconic acid | 10690 | -6.1 |
| Dodecyl sulfate | 8778 | -5.9 |
| (E)-Tetradec-1-ene-1-sulfonic acid | 6437821 | -5.9 |

**Table 4.** Vina score of test ingredients for human TMPRSS2 model

| Ingredients |  | Vina Score<br>(kcal/mol) |
| --- | --- | --- |
| Name | Pubchem CID |  |
| Inhibitor | - | -8.4 |
| Gluconic acid | 10690 | -5.8 |
| Tranexamic acid | 5526 | -5.5 |
| N-Lauroylsarcosine | 7348 | -5.4 |
| Dodecyl sulfate | 8778 | -5.2 |
| N-Lauroyl-N-methyltaurine | 61353 | -5.1 |
| (E)-Tetradec-1-ene-1-sulfonic acid | 6437821 | -5.0 |
| 6-aminocaproic acid | 564 | -4.0 |

**Figure 6.**
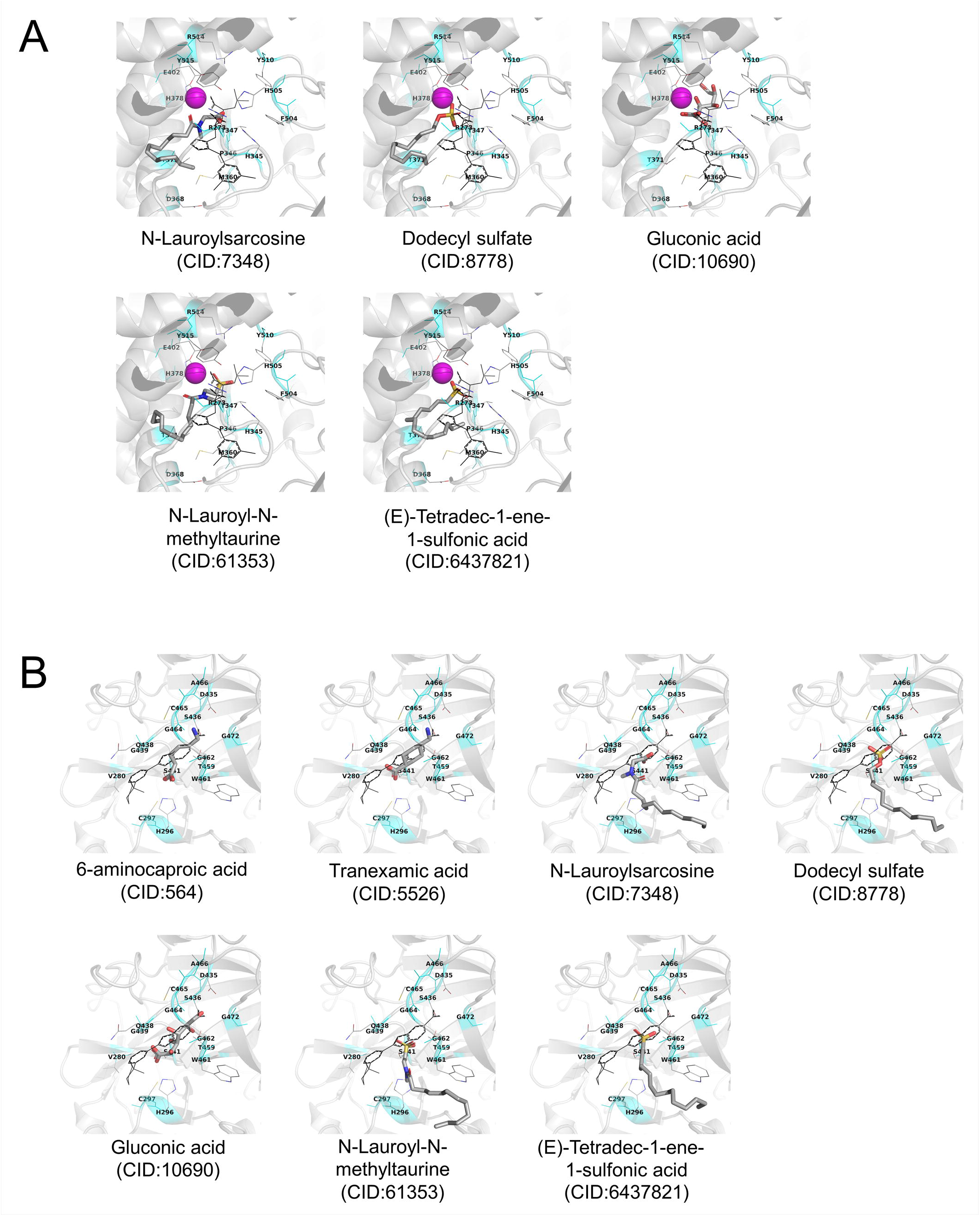
Stable docking mode of the selected ingredients obtained from AutoDock Vina docking simulations. A) Human ACE2. B) Human TMPRSS2 model. Stable docking mode of test compounds is shown as CPK-colored tubes. The original crystal structure of the inhibitor is shown in black lines. A magenta sphere shows Zn^2+^ ion. Inhibitor-binding site located at 3 Å from all heavy atoms of the crystal inhibitor is shown in cyan color. Amino acid residues located at 3 Å from the inhibitor are shown using thin tubes with label. Hydrogen atoms are neglected.

## 4 Discussion

In this study, we found that some general ingredients contained in commercially available toothpaste and mouthwash exhibit inhibitory effects on the SARS-CoV-2 spike protein-ACE2 interaction and TMPRSS2 serine protease activity, which are key factors in SARS-CoV-2 infection. To our knowledge, this is the first demonstration that toothpaste and mouthwash ingredients exhibit inhibitory effects on key host factors of SARS-CoV-2 entry into host cells.

Inhibitory effects >50% of five ingredients were observed on the SARS-CoV-2 spike protein-ACE2 interaction and seven ingredients on the TMPRSS2 serine protease activity. TDS, LMT, LSS, and SDS, which are anionic surfactants, demonstrated an inhibitory effect on SARS-CoV-2 spike protein-ACE2 interaction. For the human ACE2, these four ingredients could weakly bind to the inhibitor-binding site of human ACE2. In other words, these results suggest that these four ingredients block the interaction between the spike protein and ACE2 via binding to the inhibitor-binding site of ACE2. Since the RBD of the spike protein did not exhibit an obvious binding site for small inhibitors affecting the binding of the spike protein with ACE2, we did not investigate the docking simulations in this domain. On the contrary, surfactants, especially anions and cations, bind to proteins by electrostatic interactions, resulting in protein denaturation and interaction inhibition. Previous studies demonstrated that the RBD of the SARS-CoV-2 spike protein was positively charged at physiological pH values (the theoretical isoelectric point of RBD was 8.9) (Neufurth et al., 2020; Krebs et al., 2021). At pH 7.0-7.3 of PBS used in this study, the spike protein was presumed to be positively charged. Therefore, anionic surfactants are more likely to bind to RBD of the SARS-CoV-2 spike protein than cationic or nonionic surfactants, thereby resulting in the induction of protein denaturation and inhibitory effects. Additionally, Neufurth et al. suggested that anionic inorganic polyphosphate (poly-P) binds to the surface of RBD and prevents binding to ACE2 (Neufurth et al., 2020). These anionic surfactants and spike protein RBD interaction may result in an inhibitory effect. Based on the above results, these four anionic surfactants may act on both the spike protein and ACE2, which supports the inhibition of binding of the spike protein to ACE2. In the oral cavity, pH is maintained near neutrality (6.7-7.3) by saliva (Baliga et al., 2013), so RBD of the SARS-CoV-2 spike protein may be positively charged in the oral cavity.

SDS, TDS, LMT, and LSS also demonstrated an inhibitory effect on TMPRSS2 serine protease activity. For the human TMPRSS2 model, these four ingredients did not clearly bind to the inhibitor-binding site during the docking simulations. However, as the theoretical isoelectric point of TMPRSS2 is 8.58 (M. Pooja et al., 2021), it is presumed that TMPRSS2 is positively charged at the pH of the assay buffer used in this study. Therefore, anionic surfactants are more likely to bind to TMPRSS2 than cationic or nonionic surfactants, thereby leading to the induction of protein denaturation and inhibitory effects. Because pH is maintained near neutrality (pH.6.7-7.3) in the oral cavity (Baliga et al., 2013), the same electrostatic interaction would also occur in the oral cavity.

The GCU also exhibited inhibitory effects on the interaction between the spike protein and ACE2, as well as the TMPRSS2 serine protease activity. Docking simulations predicted that gluconic acid, which is one of parent components of GCU, could weakly bind to both inhibitor-binding sites of human ACE2 and TMPRSS2 (Tables 3 and 4). These results suggested that gluconic acid could inhibit spike protein-ACE2 interaction and TMPRSS2 serine protease activity. However, whether gluconic acid or copper ions exhibit an actual inhibitory effect is unknown. An assay for gluconic acid or copper ions alone could elucidate the mechanism of the inhibitory effect by GCU.

TXA and AHA exhibited inhibitory effects on >50% of the protease activity of TMPRSS2. TXA and AHA exhibit inhibitory effects on serine protease plasminogen (Takada et al., 1979), and in this study, researchers revealed that they also exhibit an inhibitory effect on TMPRSS2. Docking simulations predicted that TXA but not AHA could weakly bind to the inhibitor-binding sites of TMPRSS2 (Table 4). Rahman *et al* suggested that TMPRSS2 inhibitor camostat mesylate can bind to another inhibitor-binding site of TMPRSS2, whose site was estimated by MOE (Rahman et al., 2020). The possibility exists that AHA binds to another or unidentified inhibitor-binding site in TMPRSS2.

In this study, we found that seven general ingredients contained in commercially available toothpaste and mouthwash exhibited inhibitory effects on the interaction between the RBD of the spike protein and ACE2 and/or on the protease activity of TMPRSS2. Because ACE2 and TMPRSS2 are vital for SARS-CoV-2 entry into host cells, the five ingredients (SDS, TDS, LMT, LSS, and GCU) that were effective against ACE2 and TMPRSS2 may exert inhibitory effects in two steps, and a highly preventive effect on SARS-CoV-2 infection can be expected. Additionally, previous studies suggested that mouthwash ingredients, some of which were also contained in toothpaste, demonstrated antiviral properties (Carrouel et al., 2020; O’Donnell et al., 2020; Seneviratne et al., 2020). These pervious findings and our study suggest the possibility that daily oral care using toothpaste and mouthwash contribute to preventing SARS-CoV-2 infection. However, because our study involves *in vitro* and *in silico* analyses, whether SARS-CoV-2 infection can be suppressed is unclear. Therefore, we propose that an antiviral infection experiment using oral mucosa cells and upper respiratory tract cells, as well as epidemiological studies using oral care products containing these ingredients, would better clarify the antiviral activities of these ingredients and products containing them against SARS-CoV-2 infection.

## 5 Data Availability Statement

The original contributions presented in the study are included in the article further inquiries can be directed to the corresponding authors.

## 6 Author Contributions

RM and MY contributed equally and share first authorship. RM, MY, and MT conducted the research. RM, MY, TI, KT, and MT analyzed the data. RM, TI, and MT wrote the manuscript. TI, KT, SM, KK, YY, EN, and KT designed this research. All the authors read and approved the final manuscript. This paper has not been previously published elsewhere in any language and is not currently under consideration by any other publication.

## 9 Acknowledgments

The authors would like to thank Enago (https://www.enago.jp) for the English language review.

